# Dynamic RNA 3’ uridylation and guanylation during mitosis

**DOI:** 10.1101/2020.07.04.187773

**Authors:** Yusheng Liu, Hu Nie, Falong Lu

## Abstract

The successful cell division involves highly regulated transcriptional and post-transcriptional control. Some of the cell cycle related genes are periodically expressed, while most of the genes show relatively stable steady state transcript level throughout the mitotic cell cycle (Bertoli et al., 2013; Park et al., 2016). Previous TAIL-seq analysis of S phase and M phase poly(A) tail information showed that less than 2% genes showed more than 2-fold change in their poly(A) tail length (Chang et al., 2014; Park et al., 2016). In addition, the changes in poly(A) tail length between these two stages showed minimal impact on the translation of the genes as long as the poly(A) tails were longer than 20 nt. Therefore, the significance of poly(A) tail dynamics during the cell cycle remains unknown. Here, by re-analyzing the S phase and M phase TAIL-seq data, we uncovered an interesting global dynamics of RNA poly(A) tails in terms of their terminal modifications, implying global RNA regulation between mitotic cell cycles through poly(A) tail terminal modifications.

---

RNA poly(A) tail 3’ end uridylation can mark the RNA for rapid degradation (Lim et al., 2014; Morgan et al., 2017; Zuber et al., 2016). First, we looked into the uridylation frequency of RNA poly(A) tails during different stages of mitotic cell cycle. Very interestingly, the M phase RNA poly(A) tails showed much lower frequency of uridylation at its 3’ ends, especially the oligo-U, compared to that from the S phase cells as well as the asynchronized cells (Fig. S1A). Consistently, when looking into individual genes, we can see more than 90% of genes showed lower terminal uridylation frequency in M phase compared to that in S phase (Fig. S1B). We reason that the high level of uridylation in S phase may correlated with increased global RNA degradation. TAIL-seq can capture RNA both with or without poly(A) tails (Chang et al., 2014; Park et al., 2016). As poly(A) tail deadenylation is one of the most important steps in RNA degradation, we looked into the amount of RNA with deadenylated tails to estimate the RNA degradation. Interestingly, 73% of genes showed higher percentage of deadenylated poly(A) tails in S phase compared to that in M phase (Fig. S1C), confirming that higher global RNA degradation associated with uridylation in S phase. In supporting the global higher RNA degradation in S phase, we could see higher ribosome engagement of transcripts encoding core components of RNA deadenylation CCR4-NOT complexes, such as *CNOT1* and *CNOT8* in S phase than that in M phase (Fig. S1D). Consistent with uridylation of poly(A) tails deadenylated by the CCR4-NOT complex (Yi et al., 2018), *TUT4*, encoding the terminal uridylation enzyme in mammals, is also highly translated in S phase (Fig. S1E). These findings indicate that compared to M phase there is global RNA uridylation associated RNA degradation taking place in S phase.

RNA poly(A) tail 3’ end guanylation catalyzed by TENT4A/B can shield the tails from active deadenylation (Lim et al., 2018), which is opposite to the poly(A) tail uridylation. Interestingly, we could see that the guanylation in M phase cells are higher than that in S phase cells or that in asynchronized cells (Fig. S1F). When looking into individual genes, two thirds of the genes showed lower terminal guanylation frequency in M phase compared to that in S phase (Fig. S1G). This increased guanylation phenomenon in M phase may be associated with the higher RNA level of *TENT4A* (Fig. S1H). These findings indicate that compared to S phase RNA guanylation may prevent the deadenylation of the RNA in M phase.

By reanalyzing the RNA poly(A) tail information from the S phase and M phase of the mitotic cell cycle, we revealed global RNA poly(A) tail terminal modification reconfiguration during the cell cycle (Fig. S1I). The accumulation of terminal uridylation and high deadenylated transcripts at S phase suggest that there is global RNA decay happening around S phase. In turn, the accumulation of terminal guanylation and reduced deadenylated transcripts at M phase indicate that the majority of the transcriptome are protected from active deadenylation during M phase. Taken together, these data reveal that the RNA terminal guanylation and uridylation, two modifications of opposite molecular function in regulating RNA stability, are of complementary dynamics during mitotic cell cycle. Together with a recent report of m6A cell cycle dynamics (Fei et al., 2020), the findings here highlight the functional importance of RNA modifications in orchestrating the mitotic cell cycle.

## Data analyzed

This study include analysis of the following published data: TAIL-seq for asynchronized cells (GEO accession: GSE51299) and M/S phase cells (GEO accession: GSE79664), ribosome profiling data and RNA-seq data for M/S phase cells (GEO accession: GSE79664) (Chang et al., 2014; Park et al., 2016).

## Acknowledgements

This work was supported by the National Key Research and Development Program of China (2018YFA0107001) and the Strategic Priority Research Program of the Chinese Academy of Sciences (XDA24020203).

---

**Figure S1:**
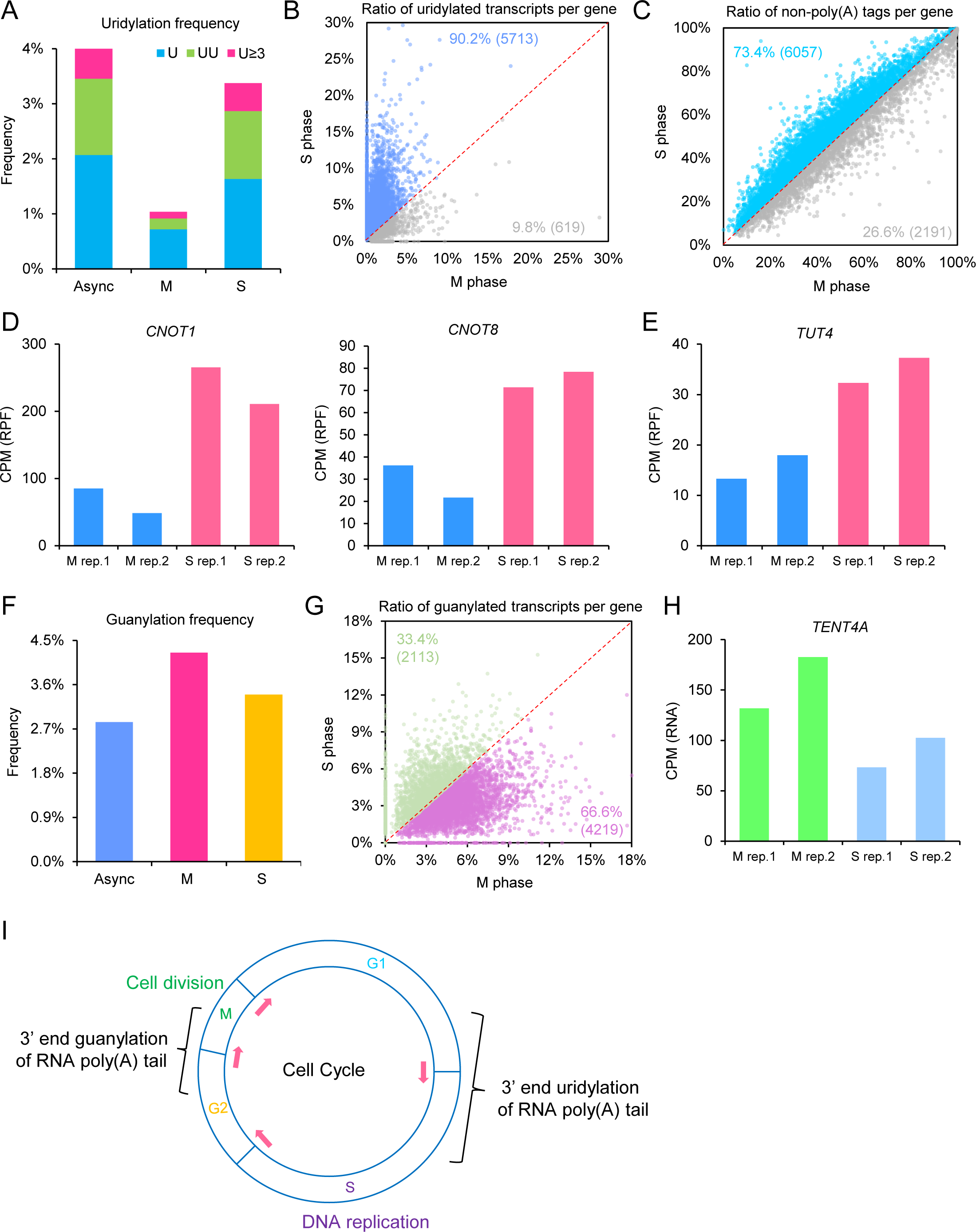
Poly(A) tail terminal uridylation and guanylation cycle during the mitotic cell cycle. (A) Frequency of uridylation is significantly lower at M phase compared to S phase. Frequency (y axis) is the fraction of detected reads with uridylation among all reads with poly(A) tails. Blue refers to mono-U, green refers to di-U, and red refers to more than 3 continuous U. Async refers to asynchronized cells, M refers to M phase cells, and S refers to S phase cells. (B) The fraction of transcripts with uridylation for each of the genes is significantly lower at M phase compared to S phase. Genes with at least 30 reads with poly(A) tails in both samples are included in the analysis. (C) The fraction of transcripts without poly(A) tails for each of the genes is significantly lower at M phase compared to S phase. Genes with at least 30 reads in both samples are included in the analysis. (D) Ribosome protected fragment counts (RPF) for *CNOT1* and *CNOT8* are higher in S phase than M phase. (E) Ribosome protected fragment counts (RPF) for *TUT4* are higher in S phase than M phase. (F) Frequency of guanylation is significantly higher at M phase compared to S phase. Frequency (y axis) is the fraction of detected reads with guanylation among all reads with poly(A) tails. (G) The fraction of transcripts with guanylation for each of the genes is significantly higher at M phase compared to S phase. Genes with at least 30 reads in both samples with poly(A) tails are included in the analysis. (H) RNA-seq quantifications of transcript level for *TENT4A* are higher in M phase than S phase. (I) A model for the transgenerational RNA cycle during mitosis.

## Notes

### Competing Interest Statement

The authors have declared no competing interest.

